# Massively parallel reporter assay–informed modeling improves prediction of context-specific enhancer–gene regulatory interactions

**DOI:** 10.64898/2026.05.01.722242

**Authors:** William DeGroat, Anat Kreimer

## Abstract

Enhancers are cis-regulatory elements that drive context-specific gene expression, yet their target genes and modes of action remain largely unresolved. Because most disease-associated variants lie in non-coding regulatory DNA, accurate, cell type–specific enhancer–gene (E–G) mapping is essential for understanding genetic risk. However, current E–G prediction frameworks lack the resolution to capture such context-specific interactions. Massively parallel reporter assays (MPRAs) provide measurement of cis-regulatory activity, but their integration into genome-scale E–G models has been limited.

Here, we introduce MPRabc, an MPRA-informed model that improves E–G interaction prediction. MPRabc integrates predicted MPRA activity, sequence-derived regulatory features, epigenomic signals, and three-dimensional chromatin contact maps with CRISPR-based perturbation training data. Benchmarking against validated regulatory interactions shows that MPRabc outperforms state-of-the-art models. We generated high-resolution E–G networks for K562, HepG2, and hiPSC cell lines and applied a graph-based framework to identify regulatory architecture, map trait-associated variants and expression quantitative trait loci, and resolve transcription factor drivers of enhancer activity. Across contexts, we accurately recovered lineage-defining regulatory programs, including GATA1::TAL1 in K562, HNF1A/B in HepG2, and POU factor circuits in hiPSCs.

Together, these results establish MPRA-informed modeling as a scalable strategy for decoding enhancer function and linking non-coding variants to gene regulatory mechanisms across cellular contexts.

## Introduction

Enhancers are cis-regulatory elements (CREs), non-coding DNA sequences that play a crucial role in tissue– and cell type-specific gene regulation [1]. These CREs contain clustered transcription factor (TF) recognition motifs that control spatiotemporal patterns of gene expression [2, 3, 4]. Although enhancers are widely recognized as hubs for disease-associated genetic variation, the mechanisms by which these CREs modulate gene expression remain partially understood [5, 6]. Identifying context-specific (e.g., tissue or cell type) enhancer–gene (E–G) links remains a critical challenge in the field, with significant implications for understanding the processes underlying complex human disorders [7, 8].

Large-scale consortia have generated, organized, and integrated extensive resources to characterize CREs across diverse cellular contexts [9, 10, 11]. Public repositories now include datasets detailing chromatin accessibility, histone modifications, TF occupancy, and three-dimensional (3D) genomic architecture [7, 9, 10, 11]. Predictive models have leveraged these measurements to score candidate E–G pairings [7, 12].

Massive strides in the training and benchmarking of E–G prediction models were made by Gschwind et al., who developed two pipelines: a systematic benchmarking workflow for E–G prediction models and ENCODE-rE2G [12]. The systematic benchmarking pipeline aggregates evidence from clustered regularly interspaced short palindromic repeats (CRISPR)-based perturbations into a gold-standard set of E–G interactions and evaluates model predictions by quantifying their ability to recover these experimentally validated links [12]. ENCODE-rE2G is a supervised machine-learning (ML) workflow that extends the Activity-by-Contact (ABC) model. The ABC model is a mathematical framework that predicts E–G pairs using a score that integrates measures of CRE activity (e.g., histone acetylation) and 3D contact frequency between CREs and candidate genes to estimate regulatory interactions [7]. ML models developed using ENCODE-rE2G are trained on CRISPR-based perturbations, providing enhanced predictive performance and interpretability through feature-importance analyses [12].

Previously, we used the ABC model to predict context-specific E–G interactions across early neuronal development [7, 8]. Next, perturbation massively parallel reporter assays (MPRAs) were applied to refine these E–G pairings further [13, 14, 15]. For perturbation MPRA, candidate enhancers were cloned upstream of a minimal promoter–reporter cassette and labeled with multiple barcodes [14]. These constructs were packaged into a lentivirus and integrated into cells [16]. RNA-to-DNA ratios measured the sequences’ context-specific activity. We evaluated wild-type and perturbed sequences to locate and characterize functional TF-binding motifs (TFBMs) [15]. We observed that CREs validated with perturbation MPRA, which demonstrated context-specific activity and interacted with TF-binding sites (TFBSs), clustered in a biologically relevant manner [8]. Based on these observations, we hypothesized that MPRA-defined activity would be an impactful feature for predicting E–G pairings.

Emerging methodologies enable more robust, context-specific analysis and interpretation of MPRA data [17, 18]. Agarwal et al. developed MPRALegNet, a deep-learning (DL) framework for predicting MPRA-based activity in three well-characterized cell lines: K562, HepG2, and WTC11 [18]. K562 is a human cell line derived from chronic myelogenous leukemia [19]. HepG2 is a human liver cancer cell line of hepatoblastoma origin and is widely used as a model for hepatocytes [20]. WTC11 is a human induced pluripotent stem cell (hiPSC) line [21]. MPRALegNet and other models with similar architectures offer a straightforward approach to featurizing MPRA experiments for E–G prediction frameworks. MPRALegNet is a high-accuracy, sequence-based model that ingests one-hot-encoded DNA sequences and outputs MPRA regulatory activity, quantified as log_2_(RNA/DNA) measurements [18]. Separate versions of the architecture are trained for each cell line [22]. Similar pre-trained DL models with distinct outputs (e.g., Sei, a framework that predicts the regulatory landscape of a DNA sequence) enable straightforward featurization by providing inputs for regulatory features without requiring genome-wide datasets or full processing pipelines [23]. This class of models enables E–G prediction models to leverage experimental information to make genome-wide predictions that would otherwise be inaccessible due to experimental data limitations. Their standardized outputs make it simple to compare regulatory feature predictions across distinct cell types or developmental contexts as well [18, 23].

Here, we introduce an MPRA-enhanced extension of the ABC framework, termed MPRabc, trained within the ENCODE-rE2G framework (Figure 1) [12]. We developed a cutting-edge E–G prediction model using features generated with MPRALegNet, Sei, the ABC model, and ENCODE-rE2G [7, 12, 18, 23]. K562 datasets were used to train MPRabc. Using the benchmarking pipeline developed by Gschwind et al., we observed performance improvements compared to existing state-of-the-art models [12]. We generated context-specific E–G pairings for the K562, HepG2, and hiPSC cell lines. Finally, we utilized the toolkit we developed, E-P-INAnalyzer, to dissect the bipartite graphs of E–G pairings and elucidate the mechanisms of E–G regulatory interactions [8, 24]; this included mapping expression quantitative trait loci (eQTLs) and single-nucleotide variants (SNVs), integrating TFBMs, and predicting regulatory substructures [8, 24]. Our approach is generalizable and can be applied to diverse cellular contexts.

**Figure 1.**
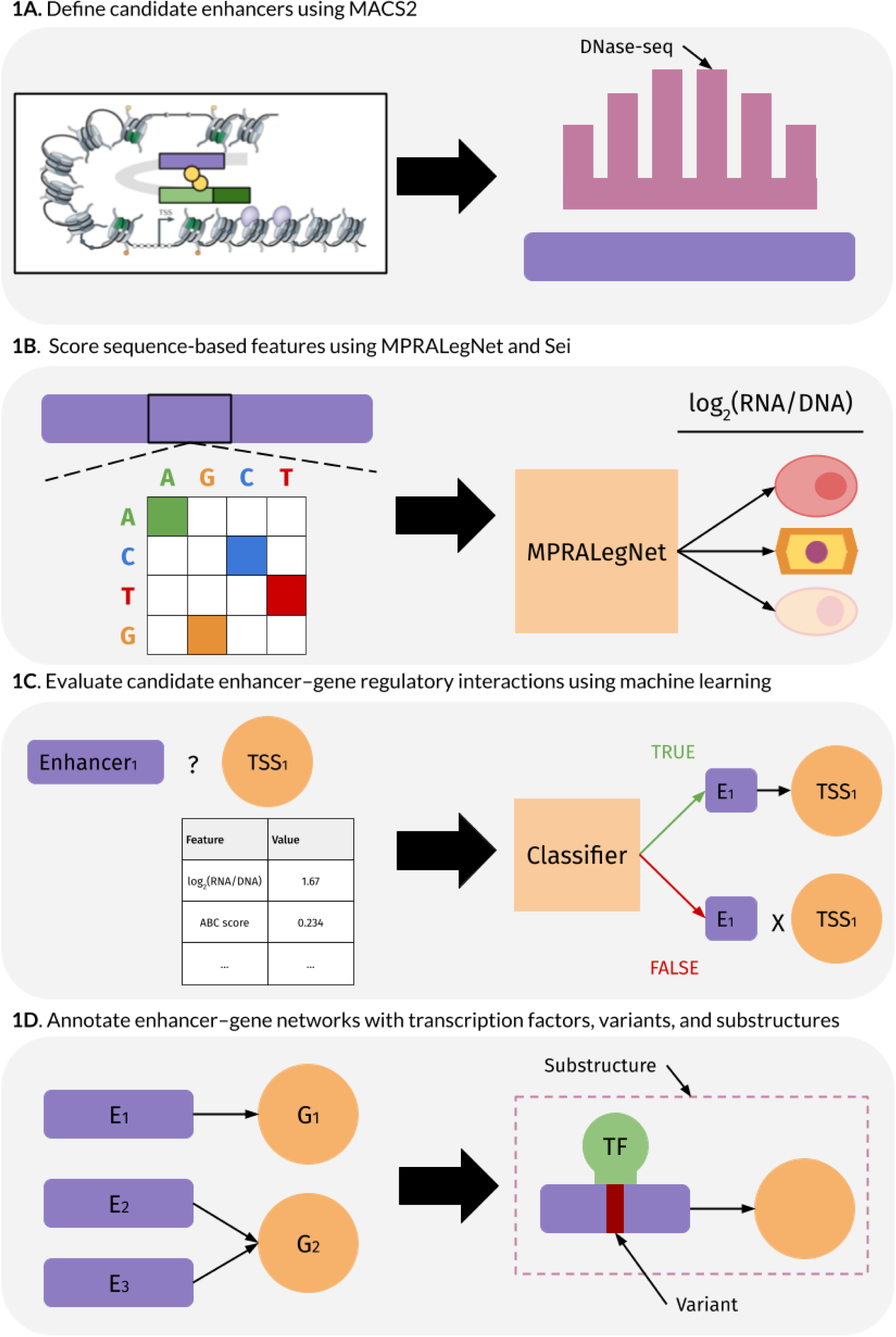
Overview of the MPRabc and E-P-INAnalyzer workflow. (A) Candidate CREs are defined from DNase-seq peaks and linked to nearby genes to create candidate E–G pairs. (B) Sequence-based features are derived by scoring each CRE with the MPRALegNet and Sei DL models. (C) A supervised ML model (MPRabc) integrates the feature set to predict true E–G interactions. (D) The resulting E–G networks are annotated with TFBMs, genetic variants, and regulatory substructures using E-P-INAnalyzer.

## Methods

### Dataset

DNase I hypersensitive site sequencing (DNase-seq), chromatin immunoprecipitation followed by sequencing (ChIP-seq) for histone H3 lysine 27 acetylation (H3K27ac), and high-throughput chromosome conformation capture (Hi-C) datasets were obtained from the Encyclopedia of DNA Elements (ENCODE) project [9]. We acquired datasets for the K562, HepG2, and hiPSC cell lines; ENCODE accessions for all datasets are listed in Supplementary 1. DNase-seq and ChIP-seq experiments were downloaded as BAM-formatted files (containing unfiltered alignments), and Hi-C experiments were downloaded as HIC-formatted files [7]. All data were available for the K562 and HepG2 cell lines. We used DNase-seq and H3K27ac ChIP-seq datasets generated in the iPS-DF19-9-11T.H hiPSC line as a proxy for the WTC11 hiPSC line [25]. Independent hiPSC lines have broadly similar chromatin accessibility and H3K27ac landscapes [26, 27]. iPS-DF19-9-11T.H and WTC11 are fibroblast-derived hiPSC lines originating from male donors, which reduces the potential confounding effects of differences in donor sex or somatic cell type of origin [21, 25]. Megamap Hi-C, an integration of Hi-C experiments across multiple tissues, was utilized for the hiPSC E–G interaction predictions [7, 12]. Both megamap and standard Hi-C versions of MPRabc are available. We defined the candidate CREs in each cell line using DNase-seq datasets (see “**ENCODE-rE2G**”).

### MPRALegNet

We accessed the MPRALegNet repository on GitHub and downloaded the pre-trained model weights from Zenodo [18, 22]. MPRALegNet is a convolutional neural network (CNN) based on the LegNet architecture [28]. The model predicts the log_2_(RNA/DNA) measurements of lentiMPRA sequences [18]. We computationally appended the MPRA adapter sequences incorporated in the MPRALegNet training constructs to each candidate CRE, ensuring that input sequences conformed to the model’s training-construct format [18]. We then applied the pre-trained model weights to each sequence to compute predicted log_2_(RNA/DNA) values [22]. In MPRabc’s current implementation, we ensemble the predictions from ten pre-trained models in the same outer test fold. We generate predictions for both forward and reverse-complement orientations and average these orientation-specific scores to obtain a single regulatory-activity score; this reduces strand-orientation bias [18]. Candidate CREs generated using the model-based analysis of ChIP-seq (MACS2) algorithm were normalized to 200 base pairs (bp) by symmetrically extending the peak from the center, matching the CRE length used for training MPRALegNet. Context-specific pre-trained models were applied for the K562, HepG2, and hiPSC cell lines. The resulting MPRALegNet-derived activity scores were subsequently used as input features in our E–G prediction pipeline.

### Sei

We downloaded the pre-trained Sei framework from the authors’ GitHub repository and applied the model without modification [23]. Sei is a deep CNN that takes 4,096-bp genomic sequences as input and predicts 21,907 epigenomic profiles across more than 1,300 cellular contexts [23]. We resized candidate CREs to 4,096-bp regions centered on the peak summit; peaks located near chromosome ends were padded with Ns to preserve the input length expected by Sei and to match the model’s training data configuration [23]. For each CRE, Sei provides probabilistic scores reflecting the likelihood that the region exhibits specific chromatin features or regulatory activity. When multiple Sei framework outputs corresponded to the same chromatin feature (e.g., multiple predictions of the same cell type–specific histone mark), we averaged the respective probabilities to obtain a single feature. The Sei-derived features used in this study are listed in Table 1, and their biological interpretation is summarized in Table 2 [29, 30, 31, 32, 33, 34, 35]. Sei does not predict CTCF, p300, or H3K4me2 features in hiPSCs. We therefore used the H1 human embryonic stem cell (hESC) line as a proxy. H1 hESCs and hiPSCs share highly similar pluripotent chromatin landscapes, making H1 a reasonable surrogate for hiPSC-specific chromatin activity [36].

**Table 1.**
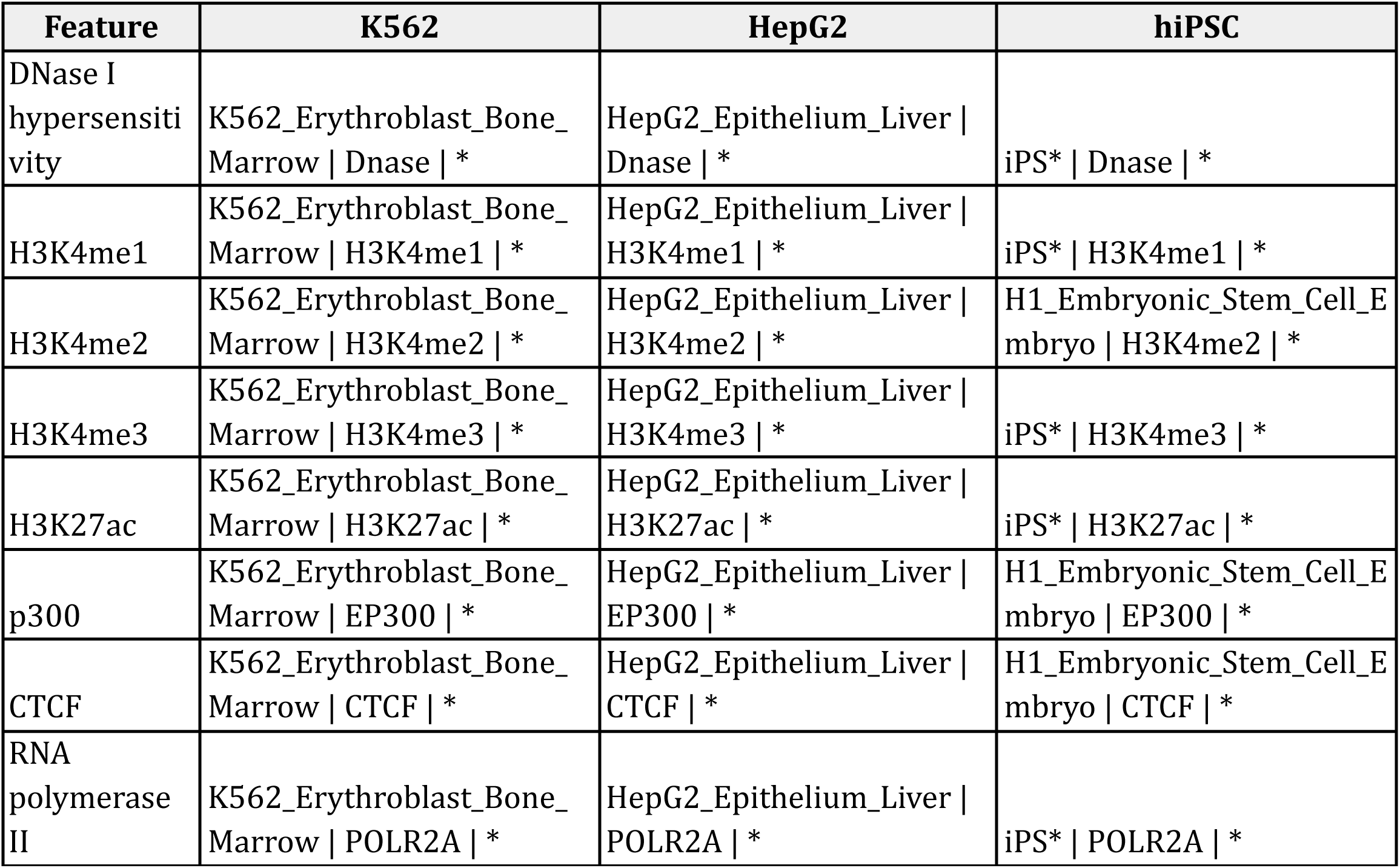
List of the regulatory features predicted by Sei and incorporated into the MPRabc feature set. For each feature, the corresponding Sei output identifiers used for the K562, HepG2, and hiPSC cell lines are provided. hiPSC features use H1 hESC proxies where Sei does not generate predictions for hiPSCs.

**Table 2.** Summary of the functional interpretation of Sei-derived features used by MPRabc. For each feature, the table provides its regulatory role and indicates whether it is characteristic of enhancers, promoters, or both.

| Feature | Description | Cis-regulatory element |
| --- | --- | --- |
| DNase I hypersensitivity | Identifies accessible, nucleosome-depleted chromatin where transcription factors can bind | Enhancer; promoter |
| H3K4me1 | Histone modification enriched at active and primed enhancers | Enhancer |
| H3K4me2 | Marks strong regulatory activity; enriched at both active enhancers and promoter-proximal regions | Enhancer; promoter |
| H3K4me3 | Sharp promoter-associated mark indicating transcription initiation | Promoter |
| H3K27ac | Marks active enhancers and promoters; distinguishes active from inactive regulatory elements | Enhancer; promoter |
| p300 | Coactivator enriched at active enhancers; correlates with enhancer activation and transcription factor occupancy | Enhancer |
| CTCF | Architectural protein defining chromatin loops and domain boundaries | Enhancer; promoter |
| RNA polymerase II | Indicates active transcription and promoter engagement | Promoter |

### ENCODE-rE2G

We integrated our sequence-based features into the ENCODE-rE2G framework, which builds on the ABC model’s approach for defining and scoring E–G interactions [7, 12]. Candidate enhancers were defined by calling peaks from DNase-seq datasets using MACS2 [37]. Peaks were resized to 500 bp, and the top 150,000 peaks ranked by total read count were retained for downstream analyses. In MPRabc, peaks were dynamically resized depending on their intended use (e.g., as MPRALegNet or Sei inputs). All gene promoters were included in the candidate set provided to the ABC model [7]. We considered all candidate CREs within a 5,000,000-bp window of each gene’s transcription start site (TSS) as potential E–G interactions. These pairings constituted the training data, allowing us to score enhancers and genes separately. Positive examples were E–G pairings validated by CRISPR interference (CRISPRi) perturbations that significantly reduced the target gene’s expression; all other candidate pairings lacking this evidence were treated as negatives (see “**Benchmarking**”) [38]. ENCODE-rE2G uses a logistic regression model to which we added L2 regularization [12]. We trained and benchmarked our model using the K562 cell line, as recommended by Gschwind et al.; the CRISPRi validation datasets were sourced from this line. We used the default feature set generated by ENCODE-rE2G from DNase-seq and H3K27ac ChIP-seq data. We added the MPRALegNet-predicted log₂(RNA/DNA) values and Sei-derived regulatory probabilities to this feature set to train MPRabc, enabling the model to leverage these new feature types alongside traditional ones [12, 18, 23]. MPRabc enabled more context-specific E–G regulatory interaction predictions by incorporating sequence-derived functional measurements of enhancer activity and sequence-intrinsic regulatory signals.

### Benchmarking

We evaluated MPRabc using the CRISPRi benchmarking pipeline developed by Gschwind et al., which harmonizes perturbation experiments into a gold-standard set of E–G interactions [12]. The pipeline assesses predictions by intersecting model scores with these labeled pairs and computing the area under the precision–recall curve (AUPRC). This is the standard metric for E–G interaction prediction model performance. The benchmarking pipeline does not perform cross-validation. Instead, models are evaluated on predefined CRISPRi datasets provided by Gschwind et al. We report performance on a dataset containing 10,356 CRISPRi perturbations (471 positive, 9,885 negative) [12]. This is the most comprehensive dataset available for any cell line [12]. Uncertainty in performance metrics was estimated using nonparametric bootstrap resampling [12]. Specifically, the pipeline generated 10,000 bootstrap replicates of labeled E–G pairs and recomputed AUPRC for each replicate. We report 95% confidence intervals (CIs) using the percentile method [12]. We adhered to the pipeline’s default settings, using the threshold corresponding to 70% recall for binary predictions. The 70% recall operating point for MPRabc corresponded to a score threshold of 0.283 (0.234 when using megamap Hi-C). To examine the contribution of individual feature groups and their relative impact on performance, we performed systematic ablations starting from the ENCODE-rE2G baseline: (1) ENCODE-rE2G, (2) MPRabc (MPRALegNet only; no Sei), (3) MPRabc (Sei only; no MPRALegNet), and (4) full MPRabc. We provide a Sei-only augmentation as a broadly applicable model, given that Sei predictions are available across >1,300 cellular contexts [23]. Because Sei-derived chromatin features (e.g., predicted DNase accessibility and H3K27ac) may be correlated with experimentally measured counterparts included in MPRabc, we quantified these relationships using both Pearson and Spearman correlation coefficients to assess redundancy and whether Sei-derived features provide a predictive signal beyond measured chromatin features. By adhering to the standardized evaluation pipeline and reporting both ablation and correlation analyses, we ensure that improvements in MPRabc can be attributed to specific feature classes and directly compared to prior E–G prediction models under consistent criteria.

### Modular community detection in enhancer-gene interaction networks using E-P-INAnalyzer

We applied our novel E-P-INAnalyzer toolkit, a generalizable framework for analyzing E–G networks, to partition regulatory substructures, contextualize trait-associated variants, and predict TFBMs in our networks of E–G pairs for the K562, HepG2, and hiPSC cell lines [8]. The predicted E–G pairs in each condition are represented as a bipartite graph [8, 24]. Nodes in this graph correspond to candidate CREs and target genes; edges are MPRabc-supported regulatory interactions. We used a multi-level modularity optimization algorithm (Louvain method) to resolve communities from the graphs. We refer to these communities as regulatory substructures [8]. We define a substructure as a community of nodes characterized by dense internal connections and sparse links to other groups, reflecting localized regulatory neighborhoods. MPRabc-generated E–G networks are highly modular; enhancers tend to regulate restricted sets of nearby genes, creating clusters with dense intracluster connectivity [7, 8]. This topology is well captured by modularity-based community detection [39]. This approach enabled us to summarize genome-scale E–G networks as collections of modules that can be annotated with additional functional information (e.g., TFBMs and trait-associated variants) [8]. Within each community, we further classified substructures according to the number of unique CREs and genes they contained, distinguishing between redundant (R) and non-redundant (NR) architectures. NR substructures are defined by the presence of a single CRE within the module (i.e., one enhancer node connected to one or more gene nodes) [8]. R substructures have multiple CREs. NR communities represent regulatory programs in which one CRE exclusively controls the target gene(s), leaving no redundant enhancer inputs. This lack of regulatory buffering renders the genes more susceptible to deleterious variants within the single CRE and creates an interpretable context in which non-coding variant–gene mechanisms can be identified without confounding contributions from additional enhancers [8, 24, 40].

### Chromosome-stratified permutation testing for trait-associated variant enrichment

To assess whether trait-associated genetic variants were enriched in condition-specific, active CREs, we performed a chromosome-stratified permutation test using all 150,000 MACS2-called DNase-seq peaks as the candidate CRE background. For each condition, we reduced the MPRabc E–G network to a unique set of CRE regions [8]. Observed overlap was defined as the number of unique variants that intersected at least one CRE in the E–G network [8]. To generate the null distribution, we preserved each CRE’s chromosome and matched each CRE to candidate regions based on its length and distance to the nearest TSS; specifically, we computed each CRE’s midpoint and assigned it to bins defined by its length and distance to the nearest annotated TSS [12]. For each permutation, we randomly sampled, with replacement, entire DNase-seq peaks from the same chromosome within matching length and distance-to-TSS bins, thereby preserving local genomic context. When no peaks were available in an exact bin, we applied a hierarchical relaxation procedure that expanded to adjacent bins until candidate peaks were identified. For each permutation, we recomputed the number of unique variants that intersected at least one CRE using the permuted CRE set. We repeated this procedure for 1,000 permutations [8]. We calculated the empirical *p*-value as the proportion of permuted values that were at least as large as the observed value, with a +1 correction in both the numerator and the denominator to avoid zero probabilities. Additionally, we report the fold enrichment (FE). Because the candidate CRE catalog includes regions that are inactive in the relevant context, enrichment within MPRabc-linked CREs provides an assessment of whether the model accurately identifies functional regulatory elements. We used fine-mapped blood trait–associated variants for the K562 cell line [19, 41]. We used liver enzyme trait–associated variants for the HepG2 cell line [20, 42]. We restricted our variant set to SNVs with *p* < 0.01. All coordinates were converted to the GRCh38 reference genome [43]. For the hiPSC line, we performed the same permutation analysis using eQTLs from the iPSC Collection for Omics Research (iPSCORE) instead of trait-associated variants [44].

### Permutation-based evaluation of concordance between predicted enhancer–gene links and eQTLs

To test whether eQTL-defined variant–gene pairs were more often supported by MPRabc-predicted E–G pairs than expected by chance, we performed a permutation-based enrichment analysis. As the observed test statistic, we quantified the number of unique variant–gene pairs in which the variant overlapped a CRE and the eQTL target gene was predicted to be regulated by that CRE. To generate a null distribution, we sampled matched sets of candidate CREs from the full DNase-seq peak catalog (see “**ENCODE-rE2G**”). For each CRE in the observed E–G network, we randomly selected, with replacement, a candidate peak from the same chromosome and class (i.e., promoter, genic enhancer, or intergenic enhancer), matched on CRE length, distance to the nearest TSS, and the number of predicted target genes [12]. This matching was performed per CRE, preserving the joint distribution of genomic and regulatory features across the sampled set. For each sampled CRE, we reconstructed predicted E–G links using the same criteria applied to the observed network, retaining edges with MPRabc scores above the threshold and restricting promoter elements to self-promoters. Each sampled CRE retained its own model-predicted target genes. This procedure preserves local genomic context and model-derived regulatory structure while randomizing the specific CREs contributing to the network. We calculated a one-sided empirical *p*-value as the proportion of permuted values that were at least as large as the observed value, with a +1 correction in both the numerator and the denominator to avoid zero probabilities. Additionally, we report the FE. eQTLs serve as a benchmark for validating predicted E–G links [12]. We downloaded whole-blood (K562) and liver (HepG2) eQTLs from the Genotype-Tissue Expression (GTEx) project [45]. We downloaded iPSC eQTLs through the iPSCORE database [44].

### Enrichment testing for cell line–specific transcription factor-binding sites

We used Find Individual Motif Occurrences (FIMO) to detect TFBSs in CRE DNA sequences from our E–G networks [46]. We downloaded 755 position weight matrices from the JASPAR CORE database and executed FIMO with a strict *p*-value threshold of 1 × 10^−5^ [47]. For each cell line, FIMO hits were restricted to CREs present in the corresponding E–G network. To compare TF-driven regulatory programs across cell lines, CREs from all E–G networks were grouped into merged genomic regions based on overlap (≥1 bp) on the same chromosome. Merged regions present in only one cell line were defined as condition-specific; merged regions present in more than one cell line were defined as condition-agnostic [8, 24]. For each TFBM, regions were scored as TFBM-present if at least one FIMO hit was detected in the region and TFBM-absent otherwise. For each cell line and each TF, enrichment in condition-specific CREs was assessed using a one-sided Fisher’s exact test on a 2 × 2 contingency table of TFBM-present versus TFBM-absent regions, testing whether TFBM-containing regions are enriched in the condition-specific set relative to the condition-agnostic set. Effect sizes were summarized as odds ratios, and we calculated pseudocount-corrected log_2_ odds ratios with a 0.5 continuity correction. We corrected for multiple comparisons across TFs within each cell line using the Benjamini–Hochberg method to control the false discovery rate (FDR) [47]. Full TFBM enrichment statistics, including raw *p*-values, FDRs, effect sizes, and region counts, are provided in Supplementary 2. This framework identifies TFs preferentially associated with cell line–specific CREs, providing evidence that the predicted E–G interaction pairings capture condition-dependent regulatory features. The enrichment of biologically relevant TFBMs within condition-specific CREs provides direct evidence that MPRabc accurately resolves regulatory drivers underlying cell line–specific E–G interactions.

### Hypergeometric survival function testing for cell line–specific gene enrichment

To assess whether genes classified as cell line–specific were overrepresented in each E–G network, we performed a hypergeometric enrichment test. We defined the background gene set as the union of all unique genes present across the K562, HepG2, and hiPSC networks (size *M*). The number of cell line–specific genes within this background was defined as *n*. For each E–G network, we then determined the total number of genes (*k*) and the subset that were condition-specific (*x*). Enrichment *p*-values were computed using the survival function of the hypergeometric distribution, **sf**(*x* − 1, *M*, *n*, *k*). This value represents the probability of observing *x* or more cell line–specific genes by chance, quantifying whether the observed representation exceeds expectations based on the background composition and network size. Statistical significance was defined as *p* < 0.05. Enrichment of cell line–specific genes in a given E–G network indicates that the MPRabc model preferentially links CREs to genes whose expression is characteristic of that cell line, supporting the biological specificity of the predicted interactions. We curated gene lists capturing transcriptional programs relevant to our three cell lines, including Myc-responsive genes for K562 (ACOSTA_PROLIFERATION_INDEPENDENT_MYC_TARGETS_UP; size = 84), liver-enriched expression for HepG2 (HSIAO_LIVER_SPECIFIC_GENES; size = 247), and pluripotency-associated features for the hiPSC line (MUELLER_PLURINET; size = 299), obtained from the Molecular Signatures Database [48, 49, 50]. Gene sets were intersected with *M* prior to enrichment analysis. The full gene lists are provided in Supplementary 3.

## Results

### MPRabc outperforms existing enhancer–gene interaction prediction models

We utilized the systematic benchmarking pipeline developed by Gschwind et al. to evaluate MPRabc (Figure 2A, Figure 2B, Figure 2C). MPRabc achieved an AUPRC of 0.707 (95% CI: 0.663–0.746), the highest performance among E–G predictors with comparable inputs (Figure 2D) [7, 12, 51, 52, 53, 54, 55, 56]. MPRabc outperformed the ABC model (AUPRC = 0.612; 95% CI: 0.567–0.656) and ENCODE-rE2G (AUPRC = 0.634; 95% CI: 0.587–0.684). At the 70% recall operating point, MPRabc achieved a precision of 0.680 (95% CI: 0.637–0.724) at a score threshold of 0.283. A version of MPRabc incorporating megamap Hi-C achieved a slightly lower AUPRC of 0.690, with a corresponding threshold of 0.234 at 70% recall. Ablation analyses (Table 3) demonstrate that both sequence-derived feature classes contribute to performance gains. Relative to the ENCODE-rE2G baseline, adding MPRALegNet-derived features increased performance to an AUPRC of 0.686 (95% CI: 0.642–0.728). Incorporating Sei-derived regulatory features further improved performance, with the full MPRabc model achieving an AUPRC of 0.707 (95% CI: 0.664–0.746). Removing Sei-derived features reduced performance, whereas removing MPRALegNet features had minimal impact on AUPRC; the Sei-only augmentation achieved performance comparable to the full model, indicating that Sei provides a broadly applicable source of predictive signal across cellular contexts. The improvement observed when adding MPRALegNet-derived features to the ENCODE-rE2G framework demonstrates that these features are biologically grounded and complementary to Sei-derived features (see “**Discussion**”). We have published the Sei-only and an MPRALegNet-only model alongside MPRabc (see “**Data and code availability**”). We quantified correlations between Sei-derived predictions and measured DNase-seq and H3K27ac signals to assess potential redundancy. Sei’s predictions showed moderate correlations with measured features (Spearman ρ = 0.48 for DNase-seq and 0.60 for H3K27ac ChIP-seq; Pearson r = 0.44 and 0.68, respectively), indicating that these features are related but not strongly collinear. Together, our findings show that incorporating MPRALegNet-defined activity and Sei-derived chromatin profile probabilities into the feature sets of E–G prediction models provides a measurable boost in accuracy beyond state-of-the-art models that rely solely on epigenomic (DNase-seq, H3K27ac ChIP-seq) and 3D genomic contact (Hi-C) datasets. K562, HepG2, and hiPSC E–G networks are available in Supplementary 4. The MPRabc model and feature table are available on GitHub (see “**Data and code availability**”).

**Figure 2.**
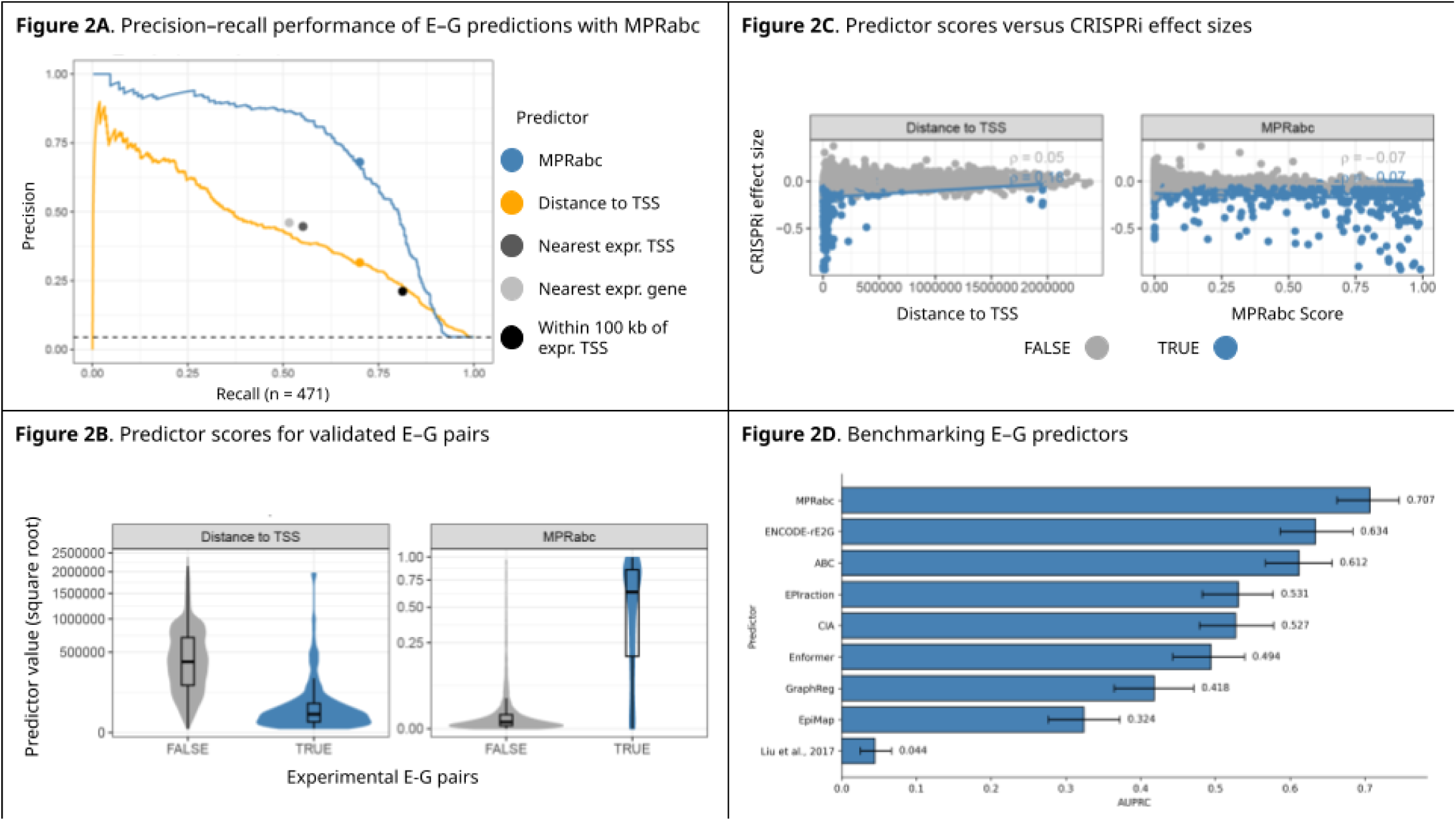
Benchmarking MPRabc on CRISPRi-validated E–G interactions. (A) Precision–recall curves for MPRabc and standard benchmarking methods on the K562 CRISPRi dataset. (B) Predictor scores versus experimental outcomes for all tested E–G pairs. (C) MPRabc scores versus CRISPRi effect size, defined as percent change in gene expression upon enhancer perturbation. Spearman correlation (ρ) quantifies the association between predictor scores and measured effect sizes. (D) Comparison of AUPRC values across E–G prediction models.

**Table 3.** Performance comparison of MPRabc and ablated variants. Performance is evaluated using AUPRC and precision at fixed recall (0.7); values are reported as means (95% bootstrap confidence intervals). Ablations remove MPRALegNet or Sei-derived features. ENCODE-rE2G is included as a baseline.

| Model | AUPRC (95% CI) | Precision at 70% Recall (95% CI) |
| --- | --- | --- |
| MPRabc | 0.707 (0.663–0.746) | 0.680 (0.637–0.724) |
| MPRabc (Sei only; no MPRAlegNet) | 0.707 (0.664–0.746) | 0.678 (0.634–0.720) |
| MPRabc (MPRAlegNet only; no Sei) | 0.686 (0.642–0.728) | 0.655 (0.610–0.695) |
| ENCODE-rE2G | 0.634 (0.587–0.684) | 0.543 (0.501–0.583) |

### K562 enhancer–gene regulatory networks are enriched for GATA1 binding sites

We utilized the E-P-INAnalyzer pipeline to investigate the E–G interaction network for the K562 cell line [8]. The network contained 48,589 unique CREs, 14,450 unique genes, and 78,023 E–G pairs [8]. The E–G network exhibited strong modular organization [8]. We applied the Louvain method to resolve communities from the network (see “**Methods**”) [8]; this revealed that 44.47% of modules were NR. Utilizing a context-matched, chromosome-stratified permutation test, we determined that blood trait–associated SNVs are enriched in CREs predicted by MPRabc to regulate a gene, compared to candidate regions (*p* = 9.9 × 10^-4^; FE = 1.814) (see “**Methods”**) (Figure 3A) [19]; this supports the hypothesis that MPRabc accurately identifies functional CREs. We tested MPRabc-predicted E–G pairs for enrichment in eQTL-defined variant–gene pairs from whole-blood GTEx data using a permutation-based approach. We observed enrichment, demonstrating that MPRabc predictions capture E–G regulatory connections consistent with eQTL-mediated gene regulation (*p* = 9.9 × 10^-4^; FE = 1.550) (Figure 3B) [45]. We predicted TFBMs in the K562 cell line using FIMO and identified TFs significantly enriched in condition-specific CREs (see “**Methods**”) [46, 47]. The most enriched TFBMs were GATA1::TAL1 (FDR = 4.8 × 10^-170^), MAFG::NFE2L1 (FDR = 6.6 × 10^-47^), and MAF::NFE2 (FDR = 1.4 × 10^-41^), consistent with the erythroid regulatory program of K562 cells, in which GATA1 and NFE2-family factors function as key regulators of differentiation (Figure 3C) [57, 58, 59]. Genes in the K562 E–G network showed significant enrichment for Myc-responsive genes (*p* = 1.8 × 10^-5^) (Figure 3D). We present an example in K562 in which a blood trait–associated SNV (chr15:31364538 C>T; rs34331965) strongly perturbs a TAL1 TFBM (alleleDiff = −0.864, motifbreakR) within an intragenic enhancer (chr15:31364317–31364817) predicted by MPRabc (score = 0.576) to regulate *KLF13* (Figure 3E) [60]. This example illustrates a plausible regulatory mechanism for the SNV rather than serving as validation of MPRabc. TAL1 is an erythroid TF, and *KLF13*, a Krüppel-like factor, has been shown to contribute to erythroid differentiation and regulation of globin gene expression, supporting the biological plausibility of this interaction [57, 61].

**Figure 3.**
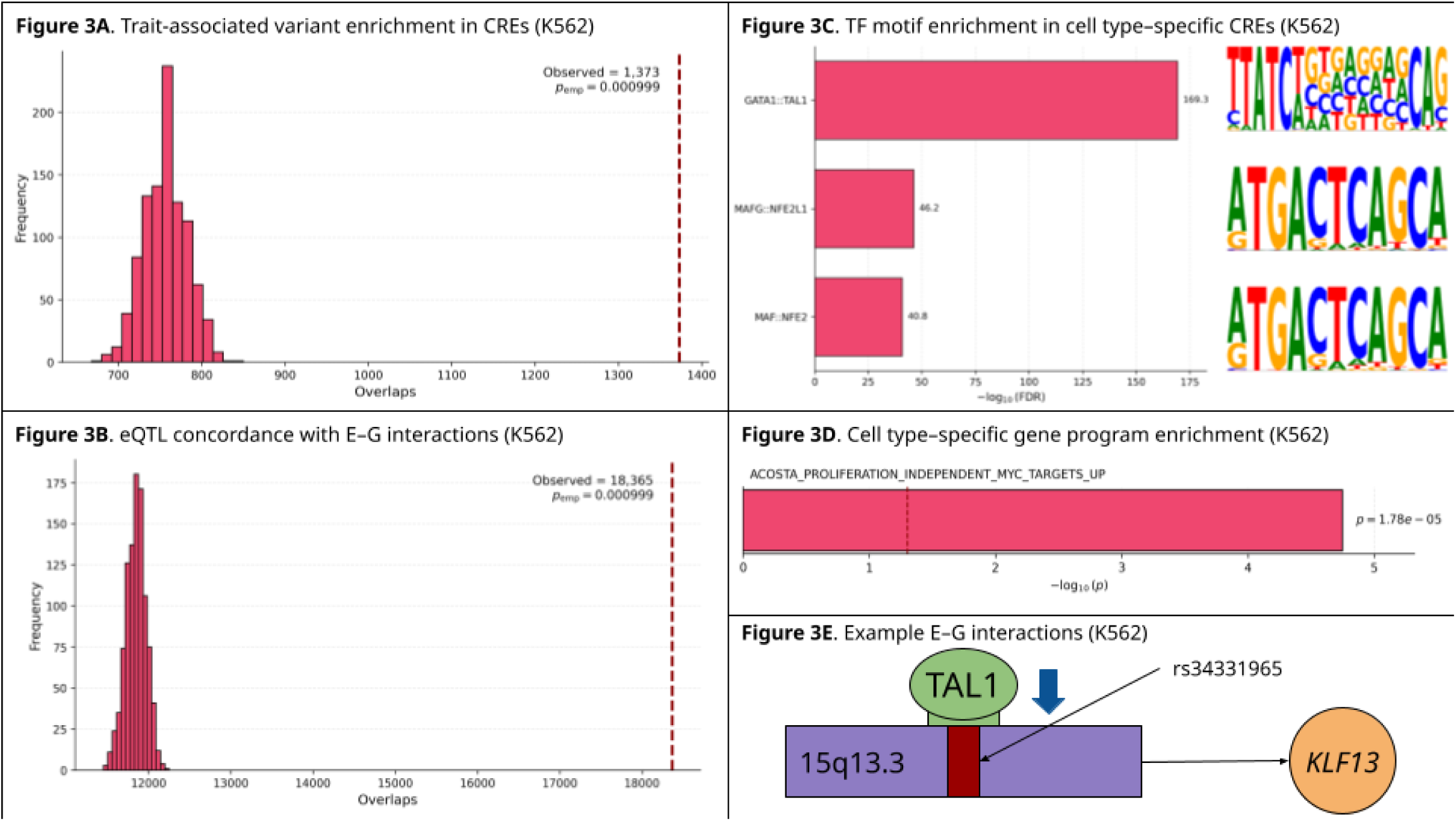
MPRabc identifies blood trait–associated regulatory architecture in K562. (A) Permutation testing shows enrichment of fine-mapped blood trait–associated variants in MPRabc-linked CREs relative to the full DNase-seq peak catalog. (B) Permutation-based concordance analysis shows enrichment of GTEx whole-blood eQTL variant–gene pairs within MPRabc-predicted E–G links. (C) TFBM enrichment in K562 CREs highlights strong enrichment for GATA1::TAL1, MAFG::NFE2L1, and MAF::NFE2. (D) Enrichment of Myc-responsive gene programs among genes in the K562 E–G network. (E) Example E–G interaction where an SNV strongly perturbs a TAL1 TFBM within an intragenic enhancer predicted by MPRabc to regulate *KLF13*.

### HepG2 enhancer–gene regulatory networks are driven by hepatocyte lineage transcription factors

We applied E-P-INAnalyzer to the HepG2 E–G interaction network, comprising 57,440 unique CREs, 14,440 genes, and 117,083 E–G pairs. Louvain community detection resolved that 53.74% of the modules were NR, indicating a high prevalence of single-CRE regulatory architectures [8]. Using a context-matched, chromosome-stratified permutation test, we observed enrichment of liver enzyme trait–associated SNVs within CREs predicted by MPRabc to regulate a gene (*p* = 9.9 × 10^-4^; FE = 1.767), demonstrating that the model accurately prioritizes functional regulatory elements in a hepatic context (Figure 4A) [20]. We next assessed the concordance between predicted E–G interactions and liver tissue eQTL variant–gene pairs [45]. Our permutation test revealed significant enrichment (*p* = 9.9 × 10^-4^; FE = 1.558) (see “**Methods**”) (Figure 4B). This indicates that the MPRabc-predicted E–G pairs capture relevant gene regulatory programs [12]. To identify the TFs driving cell line–specific gene regulation, we scanned CRE sequences using FIMO and assessed motif enrichment in condition-specific CREs. The most significantly enriched TFBMs corresponded to hepatocyte lineage-defining TFs, including *HNF4A* (FDR = 3.0 × 10⁻^161^), *HNF1A* (FDR = 8.6 × 10⁻^98^), and *HNF1B* (FDR = 4.7 × 10⁻^94^) (Figure 4C). These results are consistent with the liver-specific regulatory program of HepG2 cells, in which these factors function as core transcriptional regulators of hepatocyte identity [62, 63, 64]. Genes present in the HepG2 network showed significant overrepresentation of liver-enriched transcriptional programs (*p* = 6.6 × 10^-9^) (Figure 4D). We show an example in HepG2 in which a liver enzyme trait–associated SNV (chr13:41061165 C>G; rs17532371) strongly perturbs an HNF4A TFBM (alleleDiff = −0.773, motifbreakR) within a promoter (chr13:41060163–41062292) predicted by MPRabc (score = 0.998) to regulate *WBP4* (Figure 4E) [60]. This example presents a plausible regulatory mechanism for the SNV rather than serving as a validation of MPRabc. HNF4A is a master regulator of hepatocyte identity and liver-specific gene expression, and *WBP4*, a broadly expressed RNA-processing factor, is expressed in liver tissue, supporting the biological potential of this interaction [62, 65, 66].

**Figure 4.**
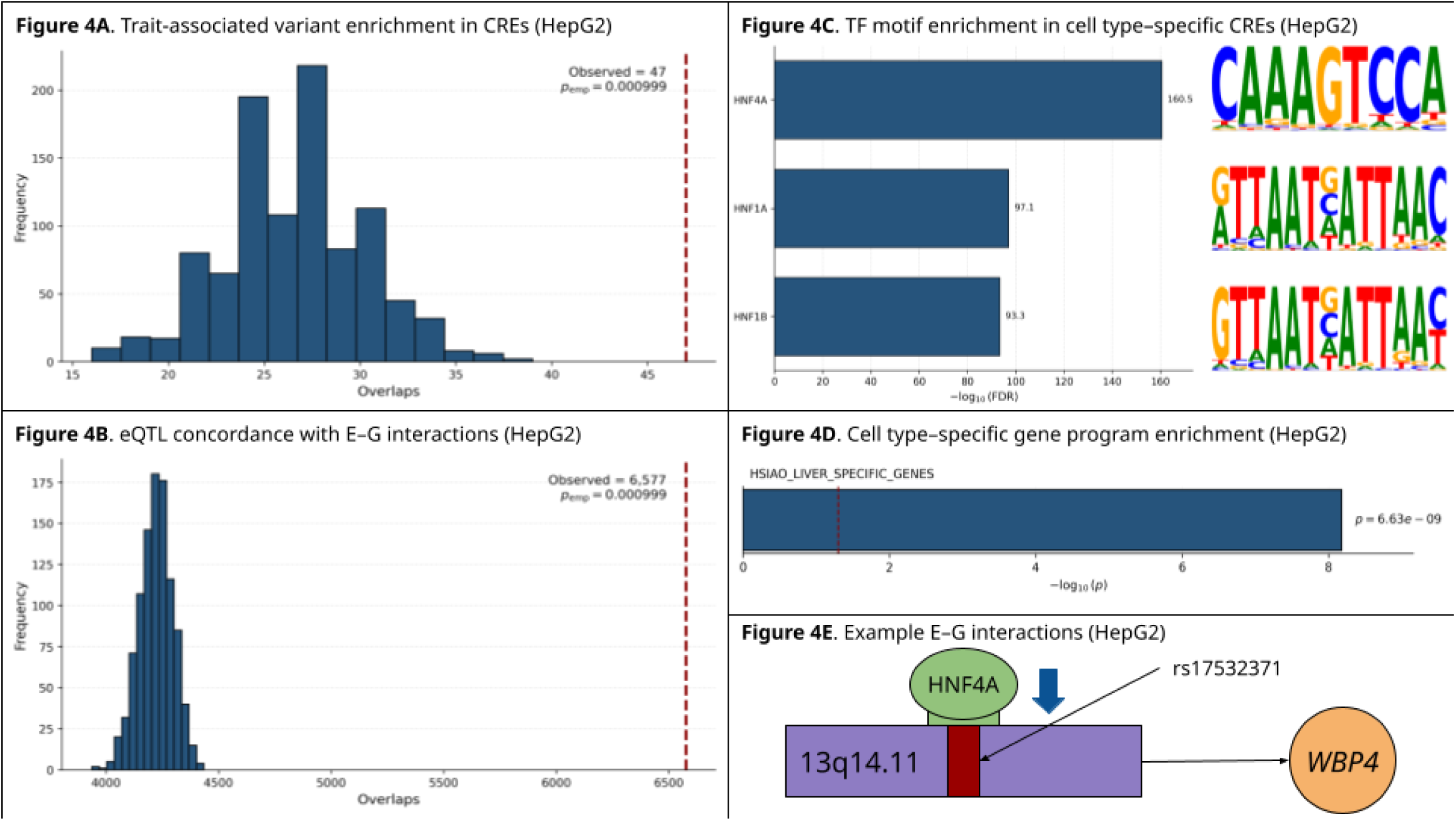
HepG2 E–G networks are driven by hepatocyte lineage transcription factors. (A) Permutation testing shows enrichment of liver enzyme trait–associated variants in MPRabc-predicted CREs. (B) Enrichment of GTEx liver eQTL variant–gene pairs among HepG2 E–G links. (C) TFBM enrichment analysis of HepG2 CREs reveals substantial overrepresentation of hepatocyte lineage-defining factors, including HNF4A, HNF1A, and HNF1B. (D) Enrichment of liver-specific transcriptional programs among genes in the HepG2 E–G network. (E) Example regulatory interaction where an SNV strongly perturbs an HNF4A TFBM within a promoter predicted by MPRabc to regulate *WBP4*.

### POU–SOX motif enrichment shapes hiPSC-specific enhancer–gene regulatory architecture

We utilized E-P-INAnalyzer to investigate the hiPSC E–G interaction network. The network contained 45,294 unique CREs, 14,436 genes, and 67,536 E–G pairs. The Louvain method resolved that 52.5% of modules were NR, indicating a substantial prevalence of single-CRE regulatory architectures [8]. Using a context-matched, chromosome-stratified permutation test, we observed significant enrichment of expression-associated variants overlapping CREs predicted by MPRabc to regulate a gene (*p* = 9.9 × 10^-4^; FE = 1.497) (see “**Methods**”) (Figure 5A). We next assessed concordance between predicted E–G interactions and variant–gene pairs defined by iPSCORE eQTLs [44]. Our permutation test revealed significant enrichment (*p* = 9.9 × 10^-4^; FE = 1.552) (Figure 5B). To determine TFs driving hiPSC-specific regulatory programs, we analyzed CRE sequences using FIMO and evaluated motif enrichment in condition-specific CREs. The most significantly enriched motifs corresponded to TFs associated with pluripotency: *POU2F1*::*SOX2* (FDR = 1.5 × 10⁻¹⁸²), *POU3F2* (FDR = 7.21 × 10^-42^), and *POU1F1* (FDR = 3.0 × 10^-38^) (Figure 5C); these findings support the well-established role of POU–SOX2 cooperative binding in maintaining the stem cell transcriptional state [67, 68]. Notably, enrichment was dominated by the composite POU2F1::SOX2 motif, highlighting SOX2-associated cooperative binding as the primary driver of the signal [69]. Genes in the hiPSC network showed significant overrepresentation of pluripotency-associated programs (*p* = 5.6 × 10^-15^) (Figure 5D). We show an example in the hiPSC line in which an eQTL (chr2:172380952 C>T; rs7596457) overlaps a POU2F1::SOX2 TFBM within an intergenic enhancer (chr2:172380665–172381165) predicted by MPRabc (score = 0.943) to regulate *ITGA6* (Figure 5E) [60]. The concordance between the predicted target gene and the gene associated with the eQTL provides independent support for this regulatory link. This example presents a plausible regulatory mechanism for the eQTL rather than serving as a validation of MPRabc. *ITGA6* encodes the laminin receptor integrin α6 (CD49f) and plays a role in stem cell adhesion and maintenance; it is expressed in hiPSCs, which supports the biological plausibility of this interaction [67, 68, 70, 71].

**Figure 5.**
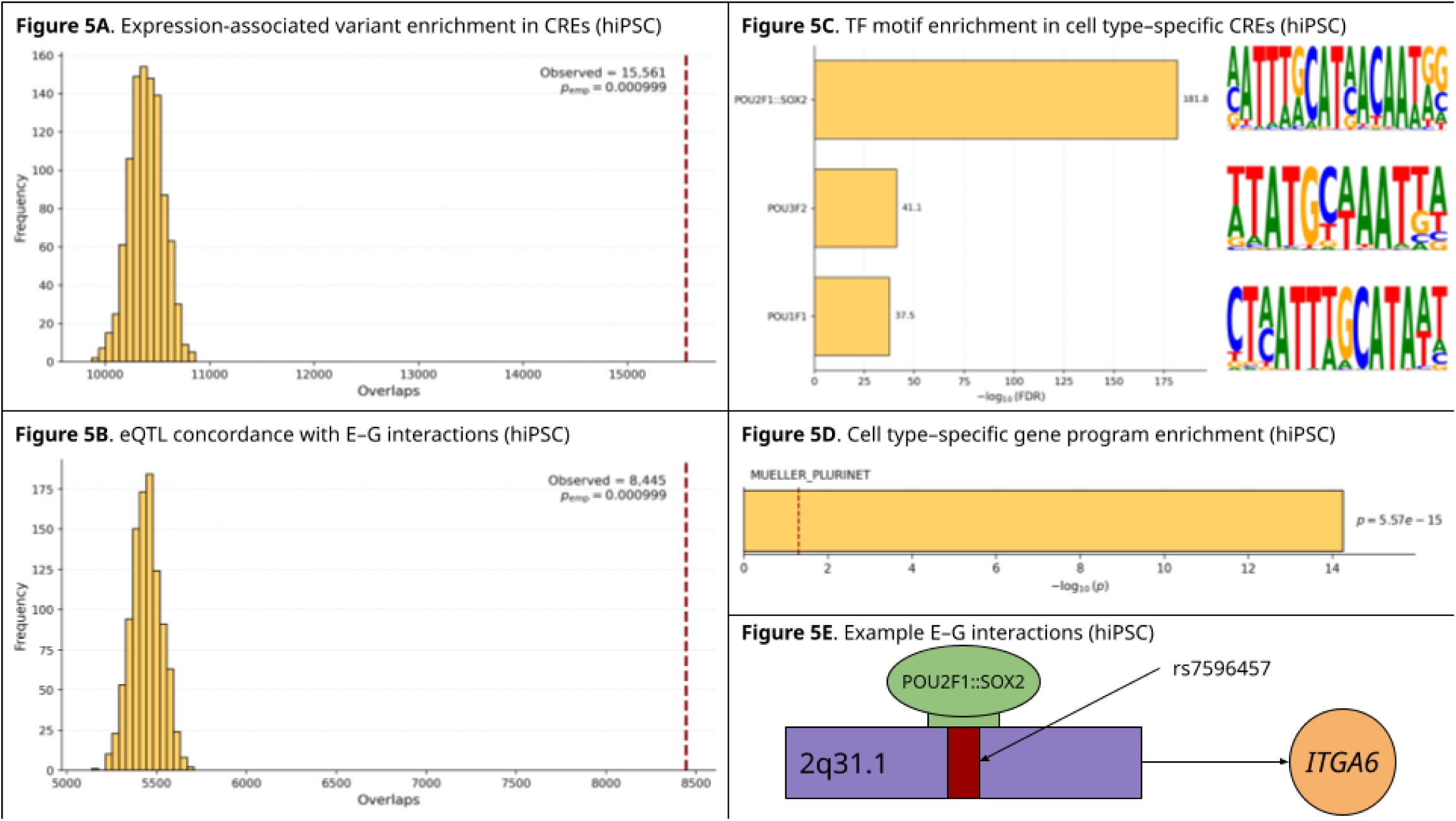
POU–SOX motif enrichment shapes E–G regulation in hiPSCs. (A) Chromosome-stratified permutation testing indicates enrichment of iPSCORE eQTL variants in MPRabc-predicted CREs active in hiPSC lines. (B) Permutation-based analysis demonstrates concordance between iPSC eQTL variant–gene pairs and MPRabc-predicted E–G links. (C) TFBM enrichment analysis identifies POU2F1::SOX2, POU3F2, and POU1F1 as the most significantly enriched TFBMs in the hiPSC E–G network. (D) Enrichment of pluripotency-specific transcriptional programs among genes in the hiPSC E–G network. (E) Example E–G interaction in which an eQTL perturbs a POU2F1::SOX2 TFBM within an intergenic enhancer predicted by MPRabc to regulate *ITGA6*.

## Discussion

Here, we demonstrate that coupling functional assay measurements with sequence-based models substantially improves CRE featurization and E–G interaction prediction [8, 12]. Using the MPRALegNet model, a CNN architecture trained on MPRA data, together with Sei, a DL framework that predicts chromatin-related sequence features, we integrated high-dimensional multi-omics and MPRA-derived functional annotations to enhance the predictive power of E–G interaction models [7, 12, 18, 23]. MPRALegNet achieved very high performance in sequence-based prediction of MPRA measurements (Pearson correlation = 0.83), approaching the assay’s technical reproducibility [18]. Sei offers an efficient, high-performance, sequence-based model for deriving diverse epigenomic features [23]. Together, these frameworks supply context-specific information that strengthens our classifier.

Benchmarking on CRISPRi-validated E–G pairs, MPRabc demonstrated superior performance [12]. The model outperformed prior models that utilized DNase-seq, H3K27ac ChIP-seq, and Hi-C datasets as their inputs [7, 12]. We note the importance of a standardized workflow for benchmarking E–G pair prediction models [12]. Integrating predicted MPRA measurements and sequence-based regulatory features within the ENCODE-rE2G framework yielded a measurable performance improvement; the MPRabc model’s AUPRC was greater than that of the best-performing implementation of the ABC model and the standard implementation of the ENCODE-rE2G model [7, 12]. These results indicate that sequence-derived regulatory features capture important context-specific signals, with MPRA-informed features providing a biologically grounded representation of regulatory activity that complements sequence-derived signals within the integrated framework.

Ablation analyses show that although Sei-derived features provide the dominant overall signal, a single MPRALegNet feature matches the contribution of multiple Sei features, indicating a compact and information-dense regulatory representation. This efficiency is particularly advantageous in less-profiled cellular contexts, where the assays underlying Sei are often unavailable but a single MPRA-informed model can provide context-specific regulatory activity.

The MPRALegNet– and Sei-derived features directly refine estimates of CRE activity, indicating that the primary improvement arises from enhanced identification of functional enhancers. However, because MPRabc is trained as a single-stage classifier over candidate E–G pairs, the contributions of enhancer activity versus target-gene assignment are not separately identifiable within this framework. Since CRE regulatory activity is inherently cell type–specific, MPRabc can generate context-specific predictions, addressing one of the principal challenges in E–G mapping [72, 73]. By pairing the MPRabc model with our previous E-P-INAnalyzer toolkit for context-specific analysis of E–G networks, we can obtain information on the architecture of regulatory programs, TFBSs, and the mechanisms underlying trait-associated variants [8, 24].

MPRabc is built on the open-source ENCODE-rE2G framework [7, 12]. We trained and benchmarked the model on K562 cell line datasets and applied it to HepG2 and hiPSC cell lines. MPRabc is engineered to be generalizable to other cellular contexts [7, 12]. The classifier does not memorize K562-specific E–G regulatory interactions. The framework is trained to capture underlying statistical regularities that govern communication between cis-regulatory elements and their target genes [7, 12]. Utilizing a supervised model with computationally engineered feature representations, including chromatin accessibility, 3D genomic contact, and predicted MPRA measurements, MPRabc learns relationships that are ubiquitous across diverse cellular contexts. Features that vary across conditions, such as chromatin accessibility, are remeasured in new contexts and substituted as inputs, enabling transfer without retraining [7]. This methodology extends across cell type–specific DL models. Incorporating outputs from a model (e.g., MPRALegNet) trained on K562 cell line data provides predictive signals that reflect sequence-encoded activity [18]. MPRabc’s architecture separates universal rules of E–G regulation from the specific data streams used to instantiate them, creating a broadly applicable framework across diverse cellular contexts.

Alternative sequence-to-activity models trained on lentivirus-based MPRA can serve as substitutes for the MPRALegNet model [18]. Currently, the MPRALegNet model is constrained to three cell lines [18]. This framework can be extended to other cellular contexts. An MPRA prediction model trained on neural contexts could be used to construct E–G networks for human cortical cells [18, 74]. This model can be used without altering the downstream classifier [12, 18]. Since the training dataset is lentivirus-based MPRA and the output is the same log_2_(RNA/DNA) measurement, a model switch preserves input–output semantics and improves context-specific sequence-based activity estimates. Large MPRA repositories, such as MPRAbase, have already been published [75]. We anticipate that sequence-based models for predicting MPRA measurements will become increasingly prevalent.

While single-cell sequencing-based approaches have advanced our understanding of E–G regulatory interaction networks, their impact is limited by data scarcity. Single-cell–based models reconstruct TF-driven regulatory programs in well-characterized cellular contexts [76]. Currently, these models are not scalable for E–G prediction across diverse datasets. MPRabc uses bulk assays in conjunction with predicted MPRA measurements. Our model’s requirements are more practical for large-scale deployments. While TF-aware modeling remains a valuable direction, our focus on data-driven CRE activity (e.g., MPRA measurements) and state-of-the-art benchmarking positions our method as a more robust and generalizable solution for mapping E–G regulatory interactions across cell types [12].

Our approach provides several advantages over large sequence-to-function models that predict E–G pairs [77]. By training directly on CRISPRi-labeled E–G pairs, MPRabc is optimized for the precise task of predicting regulatory interactions. Our model generates interpretable, auditable results with direct variant-to-consequence paths [8]. MPRabc relies on modular features, enabling us to predict E–G interaction networks for all contexts with sufficient MPRA data. This task-specific framework makes it both more practical and more directly aligned with the goals of E–G mapping.

Independent hiPSC lines (e.g., WTC11 and iPS-DF19-9-11T.H) exhibit similar chromatin accessibility and H3K27ac landscapes, but regulatory multi-omic datasets are not uniformly available for all lines [26, 27]. The absence of standardized DNase-seq and H3K27ac ChIP-seq datasets for the WTC11 cell line necessitated the use of iPS-DF19-9-11T.H as a proxy (see “**Methods**”). This may obscure subtle, line-specific regulatory features. Moreover, because Sei does not provide predictions for several key features in hiPSCs, we used H1 hESC outputs as a surrogate, thereby introducing an additional level of uncertainty [36]. These considerations should guide the interpretation of the hiPSC predictions.

Expanding the diversity of publicly available MPRA datasets will strengthen the generalizability and performance of models like MPRabc [18]. Generating libraries for new cellular contexts will provide more training data and enable transfer learning. Crucially, demonstrating the impact of MPRA data on predictive accuracy should encourage broader use of these assays. Previous studies have noted that MPRA protocols are challenging and remain largely limited to well-characterized cell lines [72]. Our findings demonstrate the added value of MPRA experiments: each new MPRA dataset can be fed into models to improve E–G interaction mapping. We anticipate that MPRAs will become a routine part of large-scale regulatory genomics projects, similar to chromatin profiling. MPRabc is an open-source, user-friendly computational model that can be adapted to encompass any cellular system, allowing for the exploration of context-specific mechanisms of gene regulation.

## Data and code availability

The MPRabc source code, pre-trained models, and scripts used to featurize the training dataset are available at https://github.com/KreimerLab/MPRabc under an MIT license. The analysis of the E–G interaction networks is available at https://github.com/KreimerLab/MPRabc-analyses, also under an MIT license. Additional datasets used in the analyses are available on Zenodo at https://doi.org/10.5281/zenodo.17822027.

## Supplementary legends

Supplementary 1. List of ENCODE accession numbers for all DNase-seq, H3K27ac ChIP-seq, and Hi-C data used in this study.

Supplementary 2. List of cell type–specific TF binding motif enrichment statistics for all TFs across K562, HepG2, and hiPSC cell lines.

Supplementary 3. Gene sets used for hypergeometric enrichment analyses of cell line–specific transcriptional programs, including Myc-responsive genes for K562, liver-specific genes for HepG2, and pluripotency-associated genes for the hiPSC line.

Supplementary 4. Table containing the E–G interaction networks for each cell line (K562, HepG2, and hiPSC). CRE coordinates, classes, target genes, and MPRabc scores are provided.

---

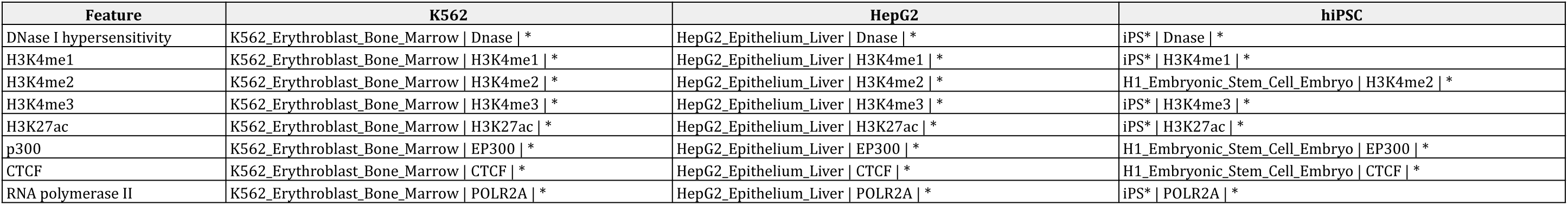

| Feature | Description | Cis-regulatory element |
| --- | --- | --- |
| DNase I hypersensitivity | Identifies accessible, nucleosome-depleted chromatin where transcription factors can bind | Enhancer; promoter |
| H3K4me1 | Histone modification enriched at active and primed enhancers | Enhancer |
| H3K4me2 | Marks strong regulatory activity; enriched at both active enhancers and promoter-proximal regions | Enhancer; promoter |
| H3K4me3 | Sharp promoter-associated mark indicating transcription initiation | Promoter |
| H3K27ac | Marks active enhancers and promoters; distinguishes active from inactive regulatory elements | Enhancer; promoter |
| p300 | Coactivator enriched at active enhancers; correlates with enhancer activation and transcription factor occupancy | Enhancer |
| CTCF | Architectural protein defining chromatin loops and domain boundaries | Enhancer; promoter |
| RNA polymerase II | Indicates active transcription and promoter engagement | Promoter |

| Model | AUPRC (95% CI) | Precision at 70% Recall (95% CI) |
| --- | --- | --- |
| MPRabc | 0.707 (0.663–0.746) | 0.680 (0.637–0.724) |
| MPRabc (Sei only; no MPRALegNet) | 0.707 (0.664–0.746) | 0.678 (0.634–0.720) |
| MPRabc (MPRALegNet only; no Sei) | 0.686 (0.642–0.728) | 0.655 (0.610–0.695) |
| ENCODE-rE2G | 0.634 (0.587–0.684) | 0.543 (0.501–0.583) |

